## Supplemental Figure legends for "Mutation in the rat interleukin 34 gene impacts macrophage development, homeostasis and inflammation in the brain and periphery"

**Supp figure 1. *Il34* and *Csf1* are broadly expressed in rat tissues and loss of IL34 has little impact on lymphoid cells and intestinal macrophages at steady-state.**

**(A)** Transcripts per million (TPM) of *Il34* and *Csf1* in tissues from the Rat atlas and juvenile rats. **(B)** Flow cytometry analysis of granulocytes in the Bone marrow, blood and spleen**. (C-E)** Single cell preparations of spleen, thymus and cervical lymph nodes (LN) from *Il34*$^{+/+}$ and *Il34*$^{-/-}$ rats were analysed for T and B cells by flow cytometry. **(F) W**hole mount imaging of small intestine (SI) in the muscularis, mucosa and villi layers from *Il34*$^{+/+}$ and *Il34*$^{-/-}$ animals carrying the Csf1r-mApple reporter. **(G)** Schematic diagram of CRISPR mediated mutation in the IL34 gene of rats; CRISPR guides are indicated in black lines and the deletion in a solid black box; solid orange blocks represent coding exons and dashed lines are introns

**Supp figure 2. Loss of IL34 does not impact the number of white matter microglia.**

**(A)** IBA1 (red or green) and DAPI (blue) were stained in brain sections from *Il34*$^{+/+}$ and *Il34*$^{-/-}$ rats. Representative images of Cerebellum, Corpus callosum and the hippocampal fissure are shown in (A). **(B)** Each dot represents counts of microglia per mm$^3$ across the different brain regions (n=3). Images were captured on the Olympus FV3000 confocal microscope and scalebars represent 100 μM.

**Supp figure 3. Loss of Il34 does not cause brain pathology.**

**(A)** Transcripts per million (TPM) derived from bulk RNA sequencing for stress induced genes (*Rbm3*, *Mt1*, *Mt2a*) and *Doublecortin* (*Dcx*) from Cortex, Hippocampus (Hipp) and Thalamus (Thal). Regions were isolated from *Il34*$^{+/+}$ and *Il34*$^{-/-}$ brains and each dot represents an independent sample sequenced. **(B)** Representative images of DCX (green) staining in the hippocampus of *Il34*$^{+/+}$ and *Il34*$^{-/-}$ animals scale bar represents 100 μM. **(C)** Morphological quantification of neuroblasts in the dentate gyrus of the hippocampus using Imaris software. Each dot is an independent animal analysed (n=3) and data is presented as area and volume stained. **(D)** Representative images of cortex stained for Glial fibrillary acidic protein (GFAP) (red) which identifies astrocytes and percentage area stained was quantified from *Il34*$^{+/+}$, *Il34*$^{+/-}$ and *Il34*$^{-/-}$ animals (n=3). **(E)** *Gfap* TPM from Cortex, Hippocampus and Thalamus. **(F)** Quantification of volume determined by diffusion imaging in the corpus callosum, anterior commissure, hippocampal commissure, inferior capsule and Diencephalon/thalamus by diffusion tensor imaging.

**Supp figure 4. Loss of grey matter microglia does not alter rat behaviour.**

Behavioural testing was performed on male **(A-C)** and female **(D-E)** rats. The sucrose preference test was performed over two 12-hour periods and consumption is represented as the weight of sucrose consumed normalized to the animal weight per testing window. **(A&D)** Sucrose preference was calculated as a fraction of volume of sucrose consumed by the total volume consumed. **(B&E)** Behaviour in the O-maze was quantified by time spent in open arms and in the open field test as time in the centre field. **(C&F)** Novel object recognition was quantified by time spent interacting with the novel and familiar objects.

**Supp figure 5. Microglia deficiency in aged IL34-deficient rats is not associated with neuropathology**

**(A)** IBA1 (red) was used to stain sections of brains isolated from animals aged 18-26 months. Representative images of cortex, hippocampus (Hipp) and thalamus (Thal). **(B)** Intact microglia were quantified in 4 fields in each brain region. **(C)** Representative images of GFAP (green) and DAPI (blue) staining in cortex, hippocampus (Hipp) and thalamus (Thal) from aged animals (18-26 months of age). **(D)** The percentage of area stained by GFAP was quantified in 4 fields per brain region. Each dot represents an independent animal (n=3). **(E&F)** *Wisteria Floribunda* agglutinin (WFA) and parvalbumin (PV) staining in the cortex from aged animals and quantification of WFA as percentage area stained, each dot represents an animal (n=3). Images were taken on an Olympus FV3000 and scale bars represent 100 μM. Statistical comparisons were calculated using a one-way ANOVA with a Tukey post-hoc test.

**Supp figure 6 Adenine-diet induced kidney injury histopathology**

**(A&B)** Cohorts of male and female rats were fed normal chow or a diet containing 0.2% adenine for 6 weeks. Kidneys from male and female animals were stained for H&E and Sirius red. Black arrows indicate adenine crystal formation. Slides were scanned with a 40x objective and digital zoom of 10x for H&E or 0.28x for Sirius red staining was used to capture images. **(C)** Lower powered (4x) images of H&E stained control and adenine treated kidneys from male and female animals.

**Supp figure 7 RNA Sequencing analysis of adenine-induced chronic kidney injury in WT and IL34-deficient rats.**

Cohorts of male were fed normal chow or a diet containing 0.2% adenine for 6 weeks and bulk RNA sequencing on kidneys was performed. **(A&B)** Genes representing the macrophage gene signature **(A)** and *Flt3* dendritic cell gene signature **(B)**. Each dot represents an independent sample sequenced (n≥4). Statistical comparisons were performed with a One way ANOVA with a Tukey post-hoc test.

**Supp figure 8 Recovery from kidney injury is not altered in IL34-deficient rats**

Cohorts of male rats were fed normal chow or a diet containing 0.2% adenine for 6 weeks followed by 4 weeks on normal chow. Control and adenine data from animals previously shown. **(A&B)** Quantification of serum urea and creatinine, **(C)** expression of *CD163* quantified by qRT-PCR and normalized to *Hprt* and **(D)** quantification of Sirius red staining area are all reduced in the recovery phase regardless of genotype. Each dot represents an independent sample (n≥4). Statistical comparisons were calculated with a two-way ANOVA with a Tukey posthoc test.

**Supp figure 9 Gating strategies for flow cytometric analysis of cell populations**

**(A)** Representative plots from analysis of blood illustrating the gating strategy used to define Monocytes (HIS48+ and CD43+), B cells, T cells and granulocytes from bone marrow, blood and spleen. **(B&C)** Representative plots of thymus **(B)** and cervical lymph nodes (cLN) **(C)** illustrating the gating strategy used to define B cells and T cells (CD4+ and CD8+). **(D)** Representative plots of brain illustrating the gating strategy used to determine Microglia and brain associated macrophages (BAM).
