## Supplemental Figures for "Mutation in the rat interleukin 34 gene impacts macrophage development, homeostasis and inflammation in the brain and periphery"

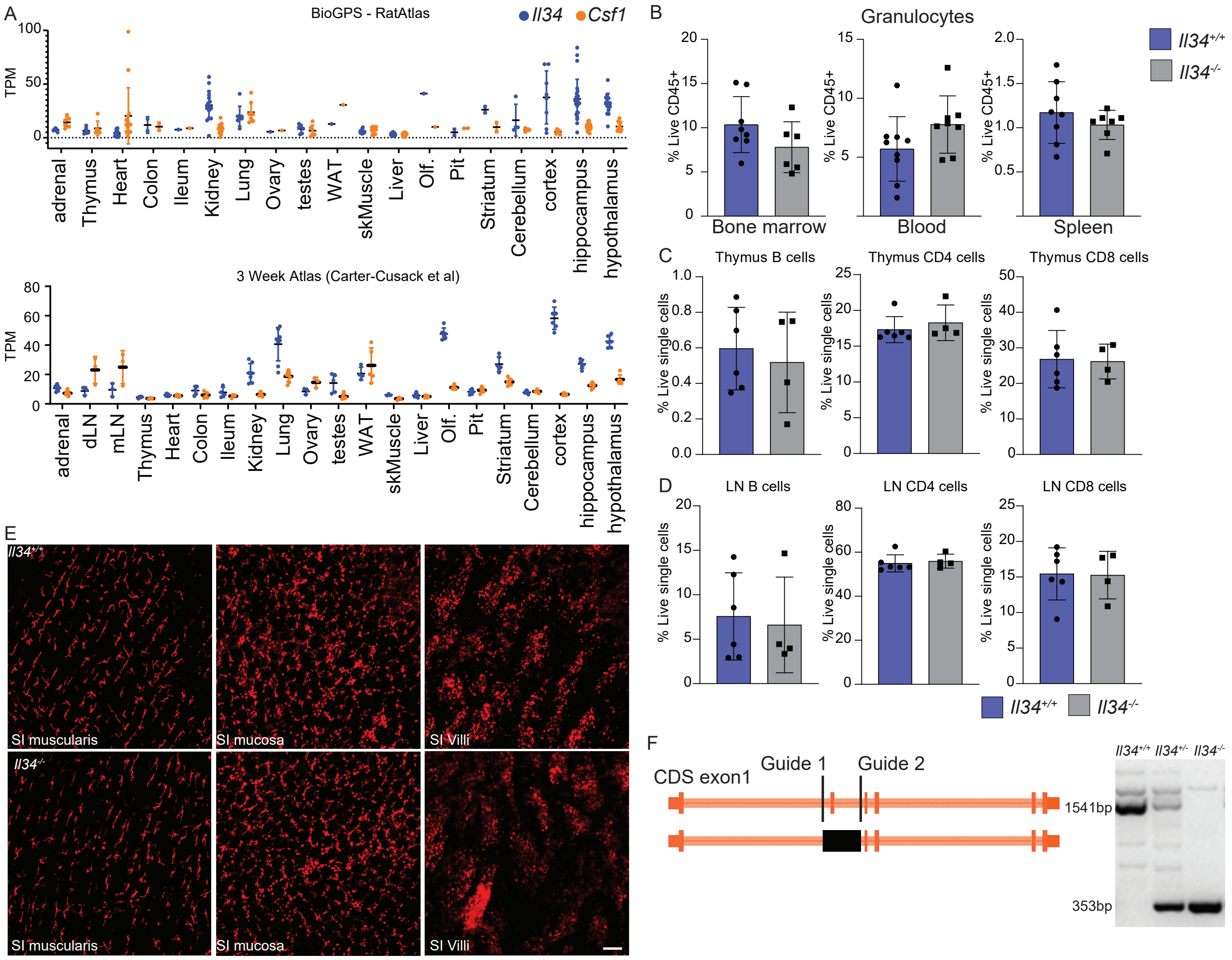


Supp Figure 1

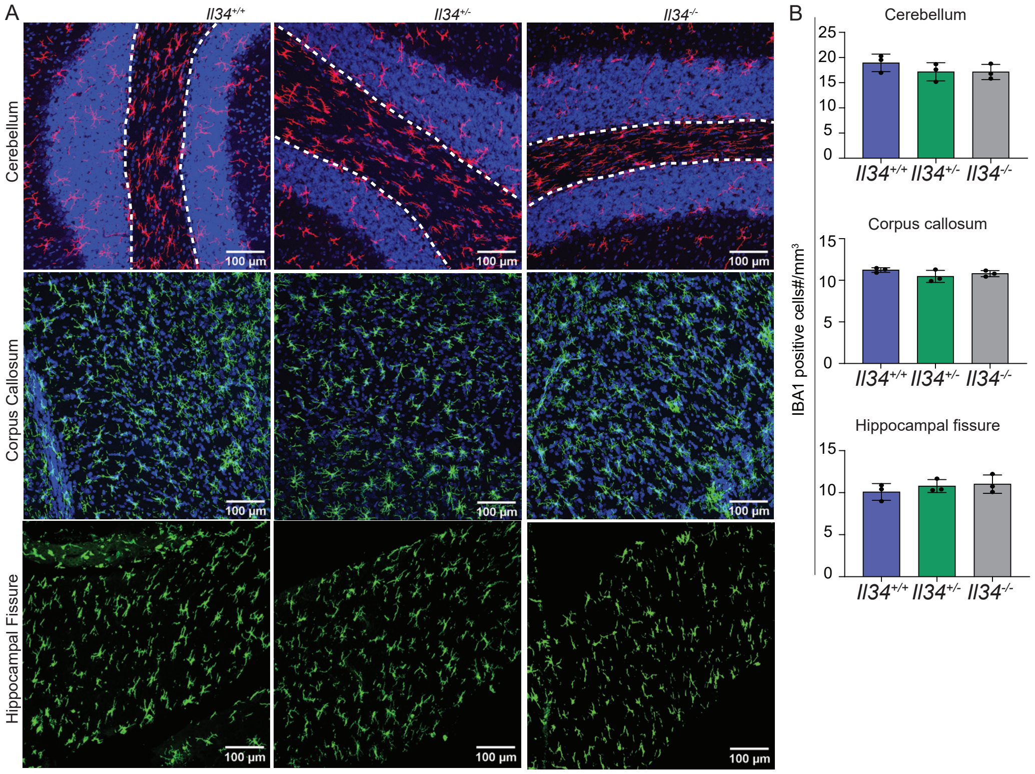


Supp Figure 2

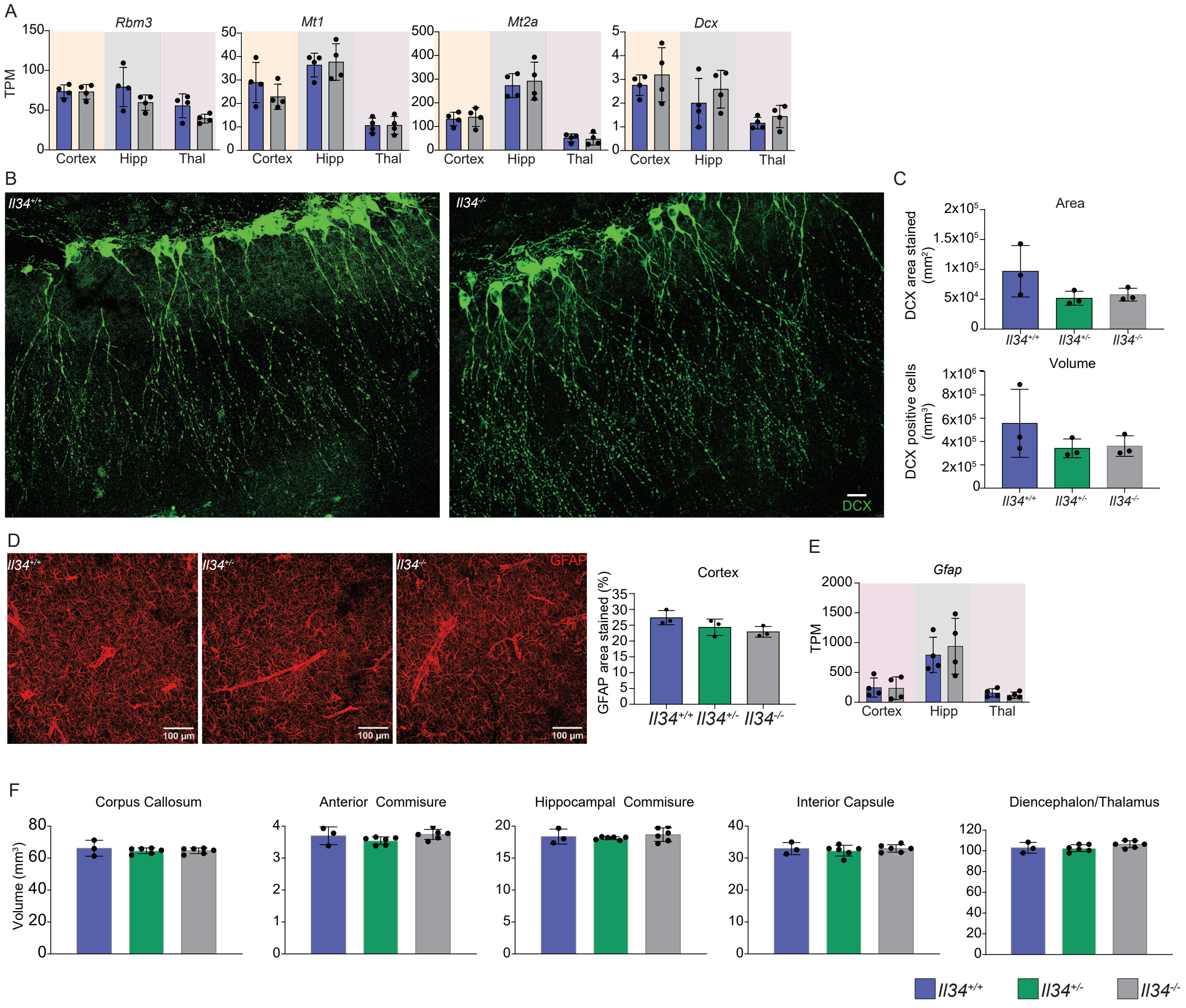


Supp Figure 3

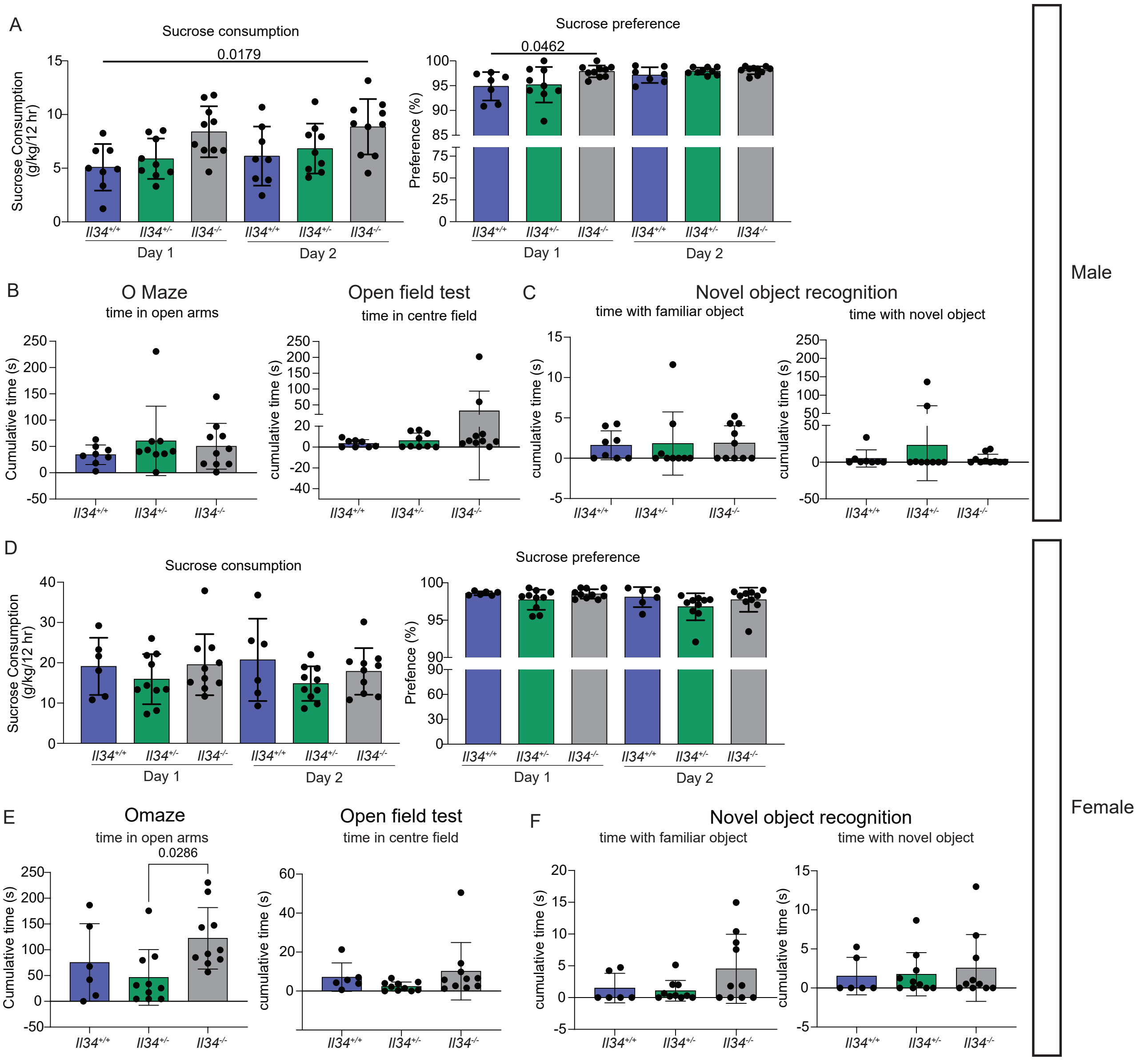


Supp Figure 4

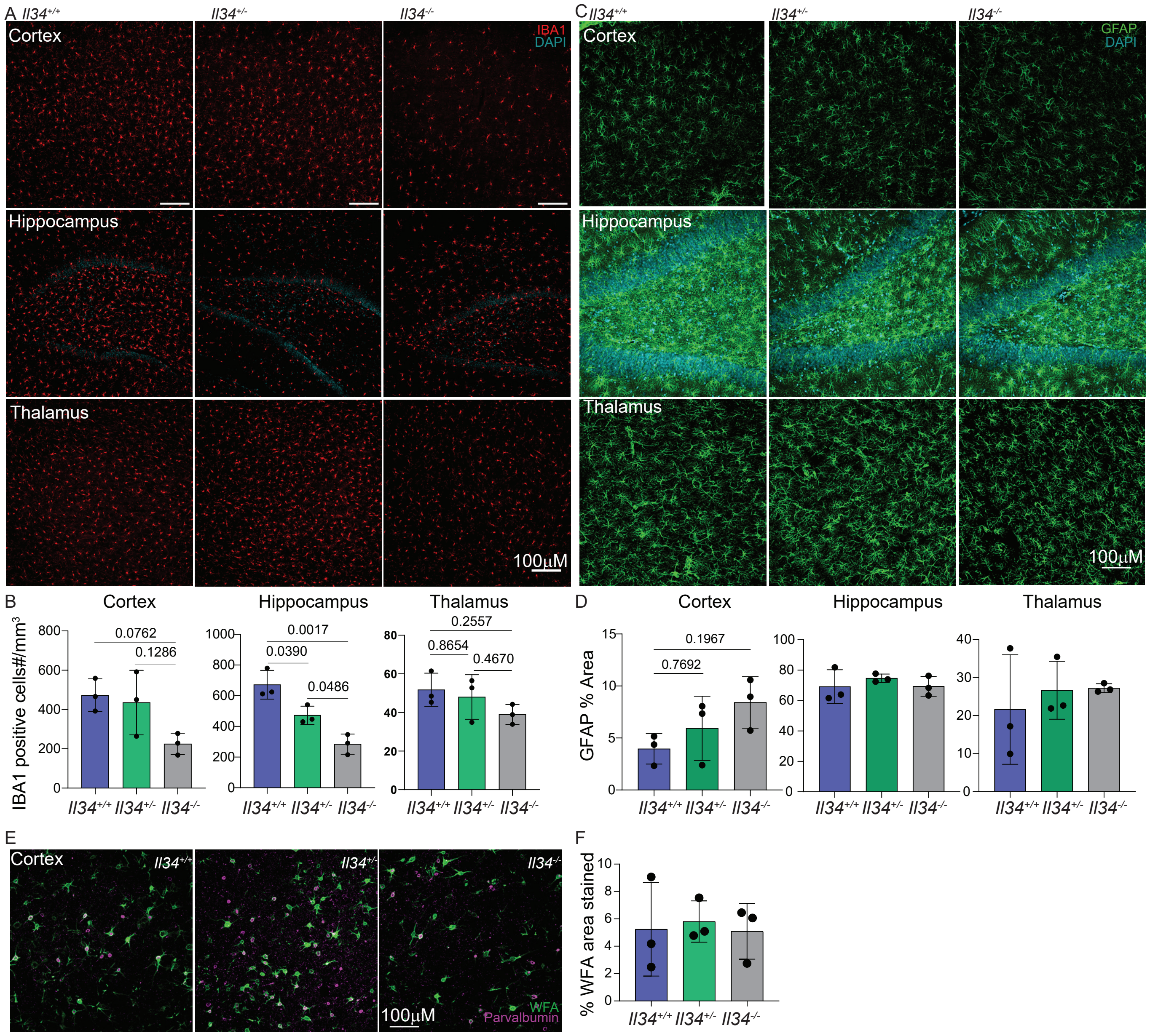


Supp Figure 5

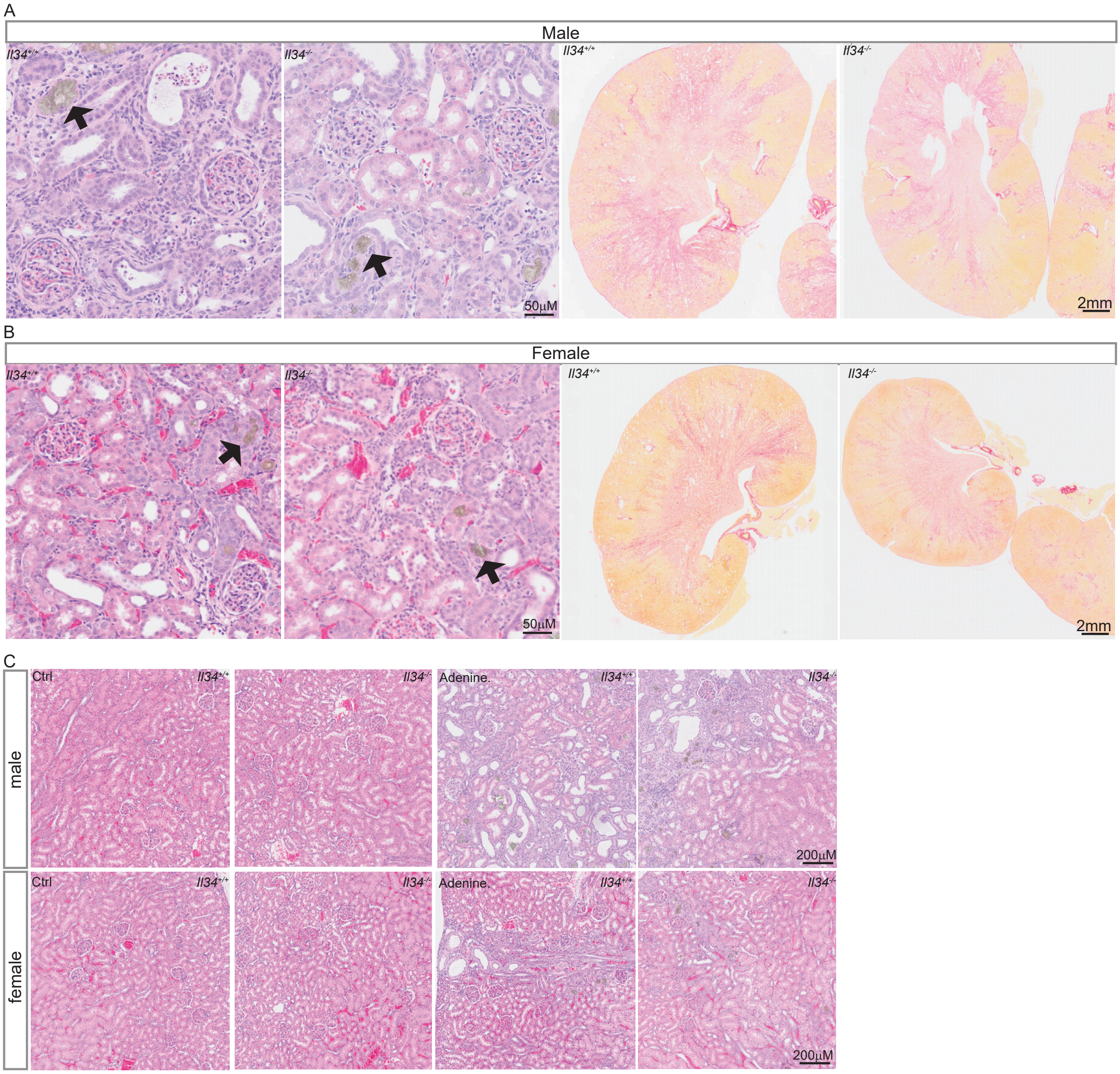


Supp Figure 6

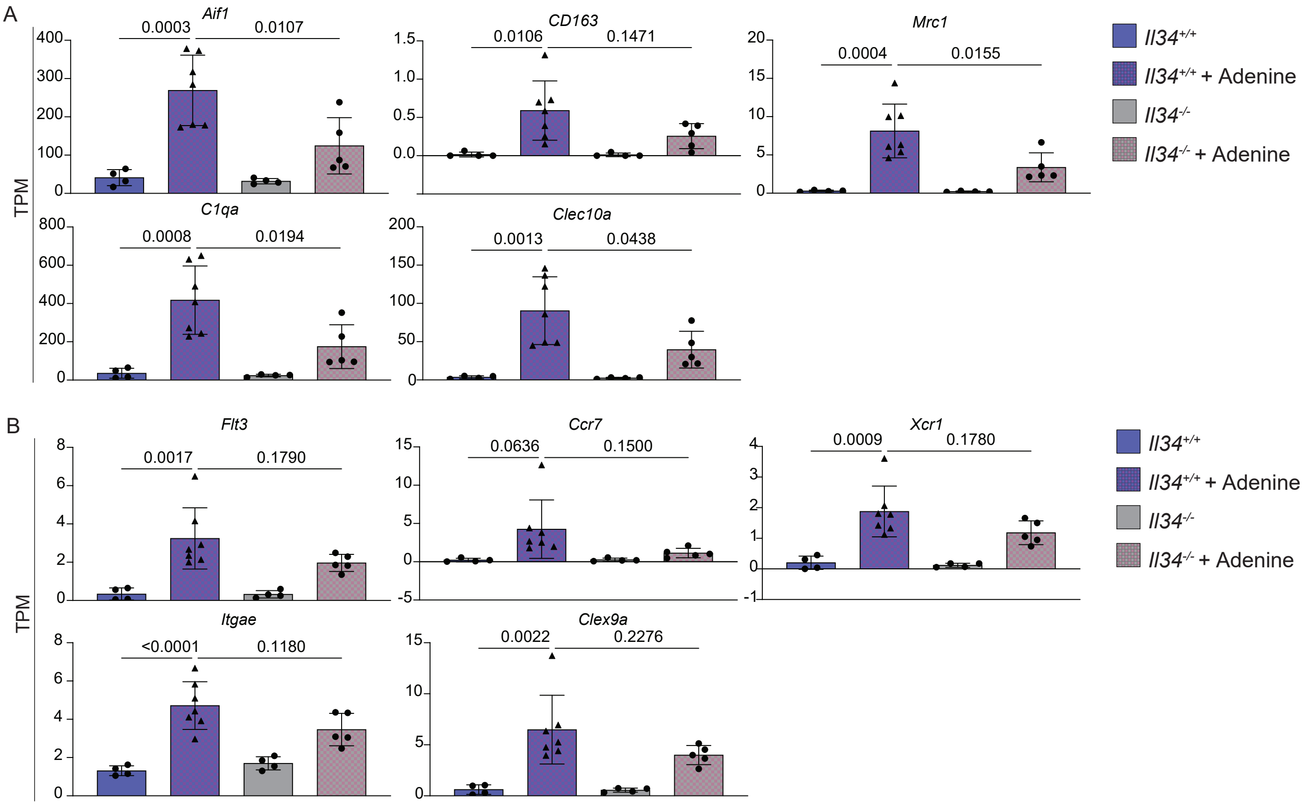


Supp Figure 7

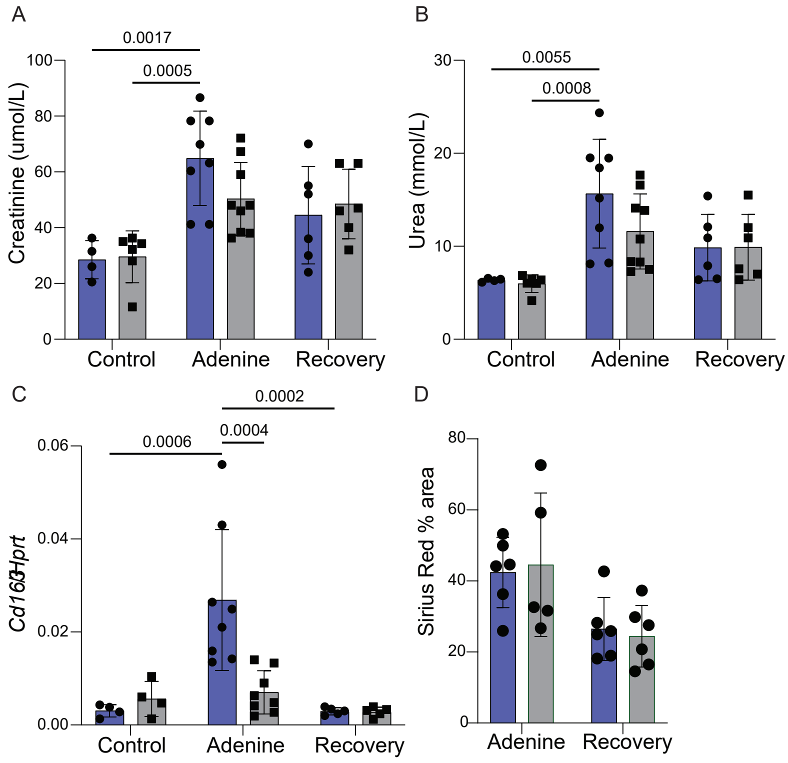


Supp Figure 8

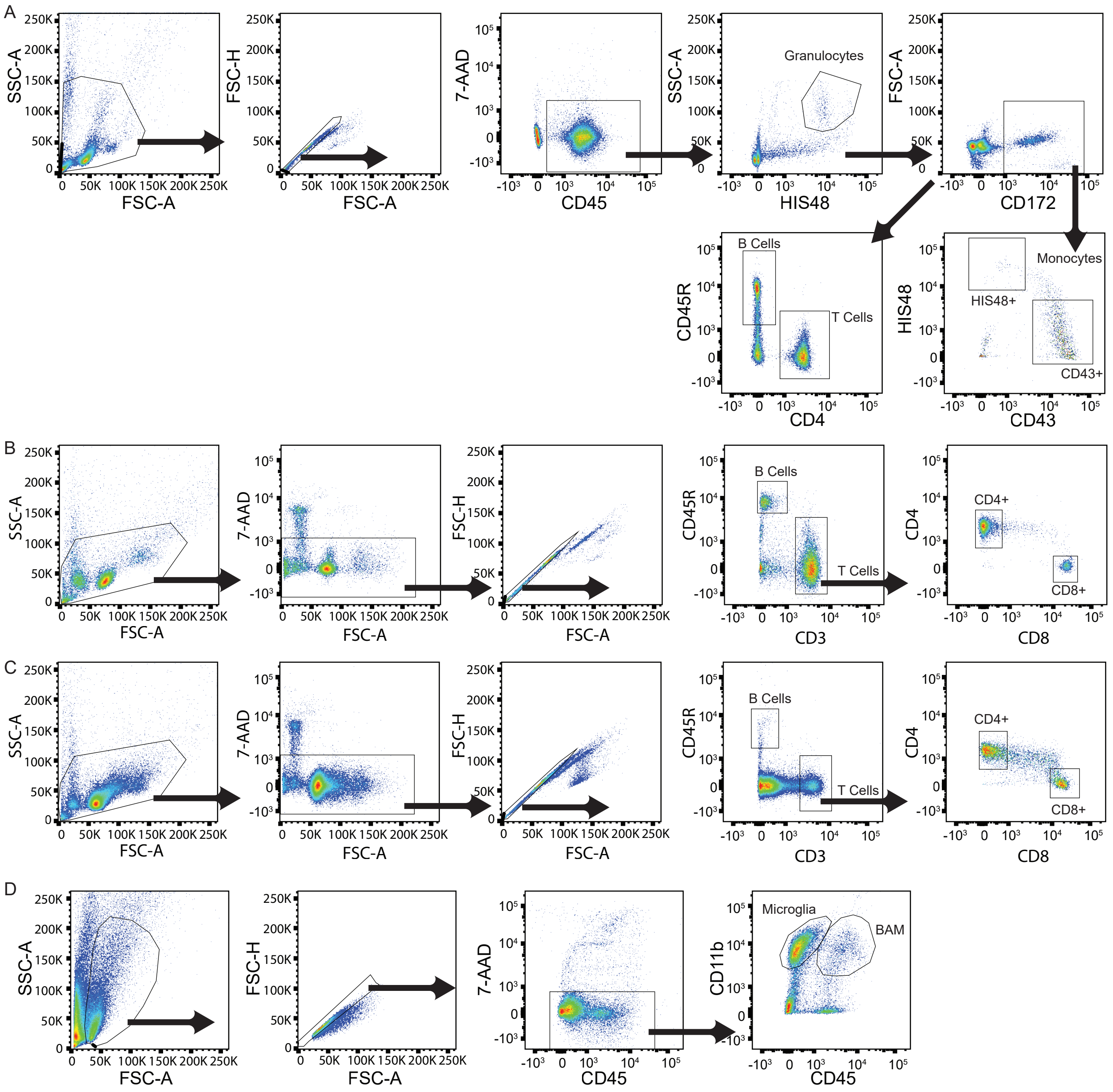


Supp Figure 9
